# Modeling the length distribution of gene conversion tracts in humans from the UK Biobank sequence data

**DOI:** 10.1101/2024.12.30.630818

**Authors:** Nobuaki Masaki, Sharon R. Browning

## Abstract

Non-crossover gene conversion is a type of meiotic recombination characterized by the non-reciprocal transfer of genetic material between homologous chromosomes. Gene conversions are thought to occur within relatively short tracts of DNA. However, the number of observable gene conversion tracts per study has so far been limited by the use of pedigree or sperm-typing data to detect gene conversion events. In this study, we propose a statistical method to model the length distribution of gene conversion tracts in humans, using nearly one million gene conversion tracts detected from the UK Biobank whole autosome data. To handle the large number of tracts, we designed a computationally efficient inferential framework. Our method further accounts for regional variation in marker density and heterozygosity across the genome, which can influence the observed length of gene conversion tracts. We allow for multiple candidate tract length distributions and select the best fitting distribution using the Akaike Information Criterion (AIC). Applying our method, we estimate that most tracts have a mean of 16.9 bp (95% CI: [16.4, 17.0]), and only a very small proportion of tracts have a much larger mean of 724.7 bp (95% CI: [720.1, 728.7]). We further estimate the proportion of gene conversion tracts with the larger mean to be 0.00525 (95% CI: [0.005, 0.00525]). After stratifying by crossover-hotspot overlap, we infer that tracts whose midpoints lie within crossover hotspots are, on average, longer than the remaining tracts.

## Introduction

During meiosis, homologous chromosomes undergo genetic recombination resulting in the transfer of genetic material. Double strand breaks that occur during recombination are resolved in two distinct ways. Crossovers result in a long tract of DNA (typically spanning millions of base pairs) being exchanged between homologous chromosomes. On the other hand, non-crossover gene conversions typically result in a non-reciprocal transfer of alleles within a short tract.^1^ These gene conversion events are thought to most commonly occur via the synthesis-dependent strand annealing mechanism, where a double stranded break is repaired by the invasion of a protruding 3’ end into the donor chromatid. Gene conversion events may also occur via the resolution of two Holliday junctions.^2^

Gene conversions can be detected in humans by analyzing sequence data from pedigrees or sperm samples and identifying positions in which the allele of one homologous chromosome has been replaced by the other.^1,3–5^ The distance between these positions, where alleles are thought to have been converted by a gene conversion event, can be used to estimate the length of the gene conversion tract. Using SNP array and whole genome sequence data from 34 three-generation pedigrees, Williams et al. determined that tract lengths are in the order of 100-1,000 bp based on detected allele conversions.^1^ Using three-generation pedigrees helps to distinguish between allele conversions and genotype errors. It can be difficult to distinguish between allele conversions and genotype errors when using two-generation pedigrees or sperm samples.

Williams et al. further identified apparent clusters of gene conversion tracts spanning 20-30 kb, which may have resulted from discontinuous gene conversion events occurring in close proximity during the same meiosis.^1^ This phenomenon has previously been referred to as complex gene conversions. Complex gene conversions as long as 100 kb were also found by Halldorsson et al.^5^ Complex gene conversions could arise from mechanisms such as GC-biased repair across long stretches of DNA.^1^ In this study, we will focus on individual gene conversion tracts where the length spanning the furthest allele converted markers does not exceed 1.5 kb.

Efforts have been made to model the length distribution of gene conversion tracts using detected gene conversion tracts in humans and other species.^6,7^ Recently, Palsson et al. detected 12,948 paternal and 15,712 maternal gene conversions transmitted to 5,420 trios in 2,132 Icelandic families.^8^ Using their model, they estimated the mean length of gene conversion tracts to be 123 bp (95% CI: [94, 135]) and 102 bp (95% CI: [71, 125]) for paternal and maternal transmissions respectively.^7,8^

Palsson et al. also found that the frequency of observed gene conversions was much higher in crossover recombination hotspots (22.4-fold and 13.7-fold for paternal and maternal transmissions respectively).^8^ While the relative frequencies of gene conversions in hotspots and non-hotspot regions have been characterized, differences in the length distribution of gene conversion tracts between these regions have not been studied in great detail.

A large number of gene conversion tracts can be detected from biobank-scale sequence data using inferred identity-by-descent (IBD) clusters. A gene conversion event occurring after the most recent common ancestor of an IBD cluster will transfer new alleles onto the haplotype, if the individual undergoing meiosis has at least one heterozygous marker within the gene conversion tract. Allele conversions cause discordant alleles within the IBD cluster in the current population, which can be used to detect past gene conversion events. Because discordant alleles can prevent the detection of the IBD cluster, Browning and Browning devised a method to use non-overlapping regions of each chromosome for detecting IBD clusters and gene conversions that have occurred on each IBD cluster.^9^ Applying their method to whole autosome sequence data from 125,361 individuals from the UK Biobank, they found 9,313,066 allele conversions inferred to belong to 5,961,128 gene conversion tracts. To detect an allele conversion, this method requires at least two haplotypes within an IBD cluster to have the same alternate allele. This means that genotype errors will not be falsely identified as allele conversions, unless the same genotype error occurs twice in the same IBD cluster.

In our study, we propose a statistical method to model the length distribution of gene conversion tracts detected from the UK Biobank whole autosome data. In our method, we account for the difference in the true length of a gene conversion tract and its observed length, which we define as the distance between the furthest allele converted markers inside this tract. The gene conversion tracts that we detect are from past transmissions in the population, for which the parental genotypes are not known. Allele conversions can only occur at heterozygous sites within a gene conversion tract in the transmitting parent, but we do not have access to the transmitting parent’s genotype data. This is not a problem in pedigree studies, where the positions of heterozygous sites in both parents are known. To appropriately account for the difference in the true and observed length of each gene conversion tract in our study without access to the transmitting parent’s genotype data, we assume that allele conversions occur with the same probability at each position within the same gene conversion tract. We estimate the allele conversion probability for each detected gene conversion tract using the heterozygosity rate of markers near the tract. Additionally, to account for the effects of linkage disequilibrium on the distribution of allele conversions, we found it necessary to exclude observed gene conversion tract lengths of one bp from our dataset, and we account for this exclusion in our analyses (see Supplementary Materials).

We allow the length distribution of gene conversion tracts to follow a geometric random variable, a sum of two geometric random variables, or a mixture of two geometric components. A geometric distribution is appropriate if the gene conversion tract grows one bp at a time, and after each extension, there is a fixed probability that it continues extending to the next bp, independent of previous steps. This distribution has been found to accurately model the length distribution of gene conversion tracts in *Drosophila*.^10^ A sum of two geometric random variables is appropriate if the gene conversion tract extends outward in both directions from a central position, with each side following the same extension process as in the geometric case. Here, we assume that the probability of extending by one bp is the same in both directions. A mixture of two geometric components is appropriate if some proportion of gene conversion tracts have a smaller mean length relative to the remaining tracts. This phenomenon has previously been observed in mammals. For example, Wall et al. estimated, applying this distribution to gene conversion tracts from a captive baboon colony, that more than 99% of all gene conversion tracts were very short (mean 24 bp), but the remaining tracts were much longer (mean 4.3 kb).^6^ Furthermore, Palsson et al. similarly estimated that within shorter gene conversion tracts (<1 kb) in both sexes, the majority of gene conversion tracts had a smaller mean compared to the remaining tracts.^8^ For each tract length distribution, we derive a closed form expression for the distribution of observed tract lengths to efficiently calculate the joint likelihood for nearly one million detected gene conversion tracts during maximum likelihood estimation. After fitting our model for each tract length distribution, we use the Akaike Information Criterion (AIC) to choose the best fitting tract length distribution.^11^

We validate our model by fitting it to detected gene conversion tracts from a coalescent simulation, originally described in Browning and Browning (2024), that incorporates evolutionary and technical factors such as mutations, genotype errors, and potential artifacts introduced by the multi-individual IBD detection method used to identify gene conversion tracts.^9^ This coalescent simulation was conducted using *msprime*, which only allows gene conversion tract lengths to be drawn from a geometric distribution.^12^ Thus, to test the robustness of our method to different tract length distributions, we run an additional simulation study drawing gene conversion tract lengths from various distributions, including a mixture of two geometric components (see Appendix).

Finally, we apply our model to estimate the mean length of gene conversion tracts detected from the UK Biobank whole autosome data. In addition to estimating the mean length for all detected tracts, we stratify detected tracts based on whether they overlap with a crossover recombination hotspot, and estimate the mean length separately for both sets of detected tracts.

## Subjects and methods

### UK Biobank whole autosome data

We ran our analysis on whole autosome sequence data from 125,361 individuals from the UK Biobank, who identified themselves as ‘white British’ in the initial release of 150,119 sequenced genomes. The UK Biobank study was reviewed and approved by the North West Research Ethics Committee and all subjects gave informed consent.^13^ The data were obtained under UK Biobank application number 19934, and the 150,119 genomes were phased using Beagle 5.4.^14,15^

### Detecting gene conversion tracts

We used gene conversion tracts previously detected in the UK Biobank whole autosome data using IBD clusters.^9^ IBD clusters are sets of haplotypes at a locus that have a recent common ancestor. If a recent gene conversion event transfers new alleles onto a haplotype in the IBD cluster, there will be discordant alleles within the IBD cluster, which can then be used to detect this gene conversion event. The detection method splits the genome into short, interleaved regions where IBD clusters are inferred or where gene conversion tracts are detected based on the inferred IBD clusters. These regions were each 9 kb long, for a total of 18 kb per IBD inference and gene conversion detection region pair, and this 18 kb pattern was repeated throughout each chromosome. Furthermore, this 18 kb pattern was offset by 0, 6, and 12 kb, and the analysis repeated for each offset to ensure that allele conversions at all positions could be detected.

Allele conversions were detected at markers where two haplotypes in an IBD cluster shared one allele and two others shared the alternative allele, minimizing the false detection of genotype errors as allele conversions. Furthermore, only markers with minor allele frequency (MAF) ≥ 5% were considered to avoid misclassifying mutations as allele conversions.

After allele conversions were detected, they were clustered to form detected gene conversion tracts. Allele conversions were considered to belong to the same gene conversion tract if they were located within 1.5 kb of each other, and if the membership of the two sub-clusters (representing the two alleles present in the IBD cluster) overlapped for the two allele conversions.

Across all the autosomes, 9,313,066 allele conversions were detected.^9^ These allele conversions were inferred to belong to 5,961,128 detected gene conversion tracts. Furthermore, 4,943,183 (82.9%) of the detected gene conversion tracts were comprised of a single allele conversion.^9^ 1,017,945 (17.1%) of the detected tracts were comprised of two or more allele conversions. We refer to the length spanning the furthest allele converted markers in a detected gene conversion tract as the observed tract length of the gene conversion tract. If a detected gene conversion tract is comprised of a single allele conversion, the observed tract length is one bp.

We label the observed tract lengths of all detected gene conversion tracts as {ℓ_*j*_|*j* = 1, …, *m*}. The procedure used to detect gene conversion tracts in each offset assumes that gene conversion tract lengths do not exceed 1.5 kb. To take this into account, we exclude any observed tract lengths exceeding 1.5 kb when estimating the mean gene conversion tract length. This results in the exclusion of 141,361 tracts (2.4% of all detected tracts). We also exclude observed tract lengths of one bp prior to estimation, because our model assigns a higher probability mass at one bp compared to what we observe in the data (see Supplementary Materials). This is likely because we do not account for linkage disequilibrium in our model. Although we exclude observed tract lengths of one bp when estimating the mean gene conversion tract length, the proportion of observed tract lengths of one bp is used to understand the effect of linkage disequilibrium on the distribution of observed tract lengths (see Supplementary Materials). We appropriately account for the omission of these tracts in our model by truncating the marginal distribution of observed tract lengths (derived in a later section) at one bp and 1.5 kb. After removing both detected tracts of 1 bp and those exceeding 1.5 kb, we are left with 876,584 detected tracts. Although excluding these tracts reduces the amount of data used in the estimation procedure, results from our simulation study suggest that the resulting estimates are unbiased under the truncated model.

### Definitions and overview of model

We model *N*, the length of a gene conversion tract, as a geometric random variable, a sum of two independent and identically distributed geometric random variables, or a mixture of two geometric components. We further let *L* be a random variable representing the observed tract length of a gene conversion tract, which is the length spanning the furthest allele converted markers within the gene conversion tract. The event *L* = 0 represents no allele conversions occurring within the tract, and *L* = 1 represents one allele conversion occurring within the tract. In the following sections, we derive the conditional distribution of *L* given *N* and the marginal distribution of *L*. We further describe the procedure we use to obtain a maximum likelihood estimate of 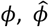, using the observed tract lengths {ℓ_*j*_|*j* = 1, …, *m*} detected from the UK Biobank whole autosome data.

### The distribution of the observed tract length conditional on the gene conversion tract length

The observed tract length of a gene conversion tract, represented by the random variable *L*, depends on where allele conversions occur on the gene conversion tract. We will first assume that allele conversions happen with probability *ψ* at every position within some gene conversion tract that is exactly *n* bp long. Under this scenario, the following conditional distribution has previously been derived.^16^

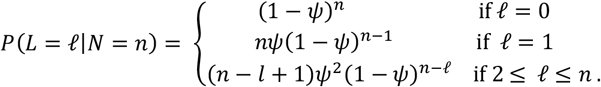

In the probability above, we conditioned on the gene conversion tract length, represented by the random variable *N*, being *n* bp long. Obtaining an observed tract length of zero bp is equivalent to allele conversions not occurring within the gene conversion tract, which happens with a probability of (1 − *ψ*)^*n*^. Next, obtaining an observed tract length of one bp is equivalent to an allele conversion occurring at exactly one position within the gene conversion tract. There are *n* possible positions in which the allele conversion can occur, and each configuration happens with a probability of *ψ*(1 − *ψ*)^*n*−1^. Lastly, to obtain an observed tract length of ℓ bp, where 2 ≤ ℓ ≤ *n*, we need to observe two allele conversions that span exactly ℓ positions, and allele conversions cannot occur at the *n* − ℓ positions flanking the two allele conversions. There are *n* − ℓ + 1 ways to overlay these two allele conversions on the gene conversion tract, and each configuration occurs with a probability of *ψ*^2^(1 − *ψ*)^*n*−ℓ^.

### Deriving the marginal distribution of the observed tract length

If the gene conversion tract length *N* is drawn from geometric distribution with mean *ϕ*, we have,

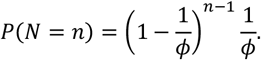

Letting λ = 1/*ϕ*,

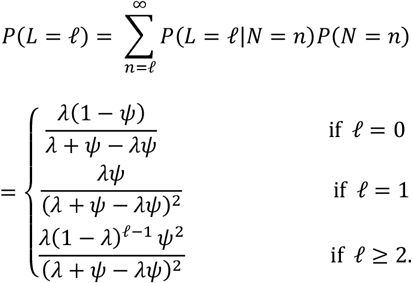

This is the marginal distribution of the observed tract length *L*. A closed form expression for *L* was not derived previously, but this form is crucial for accelerating likelihood computations, given that we compute the joint likelihood of nearly one million observed tract lengths during maximum likelihood estimation. We further truncate this distribution to appropriately model observed tract lengths detected in the UK Biobank sequence data using the multi-individual IBD method.^9^ Recall that we only retain observed tract lengths between 2 and 1,500 bp during estimation, so we account for this by truncating the distribution of *L* between 2 and 1,500 bp.

We have,

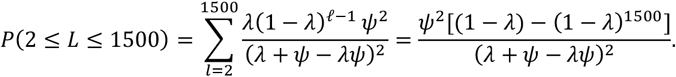

Then,

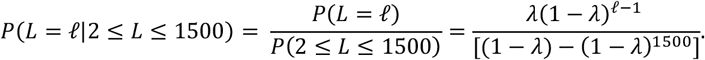

Notice that conditioning on 2 ≤ *L* ≤ 1500 removed the parameter *ψ* from our model.

As mentioned earlier, {ℓ_*j*_|*j* = 1, …, *m*} represents the observed tract lengths in our dataset. When fitting the model, we use the filtered set of observed tract lengths, {ℓ_*j*_|*j* = 1, …, *m*, 2 ≤ ℓ_*j*_ ≤ 1500}. Henceforth, we will also index our random variable *L* using *j. L*_*j*_ represents the random variable corresponding to the observed tract length of detected gene conversion tract *j* in our dataset. We have,

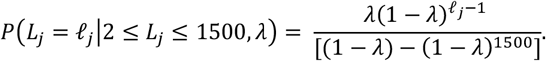

We also consider two alternative distributions for *N*: a sum of two independent and identically distributed geometric random variables, and a mixture of two geometric components. The derivations of *P*(*L*_*j*_ = ℓ_*j*_|2 ≤ *L*_*j*_ ≤ 1500) under both settings are provided in the Appendix. Under these settings, *P*(*L*_*j*_ = ℓ_*j*_|2 ≤ *L*_*j*_ ≤ 1500) depends on *ψ*_*j*_, so we estimate *ψ*_*j*_ for each tract *j* before estimating *ϕ*. The procedure to estimate *ψ*_*j*_ for each tract *j* is described in the following section.

### Estimating the allele conversion probability for each detected tract

Recall that *ψ*_*j*_ represents the probability that an allele conversion will occur at each position within detected gene conversion tract *j*. When *N* is a sum of two geometric random variables or a mixture of two geometric components, the likelihood of the observed tract length for detected gene conversion tract *j, P*(*L*_*j*_ = ℓ_*j*_|2 ≤ *L*_*j*_ ≤ 1500), depends on *ψ*_*j*_ (see Appendix), so we need to estimate *ψ*_*j*_ for *j* = 1, …, *m* to obtain a maximum likelihood estimate for the mean gene conversion tract length *ϕ*.

Allele conversions occur at positions within each gene conversion tract where the individual is heterozygous. Therefore, the probability that a randomly selected individual from the population is heterozygous at a given marker can be used to estimate the probability that an allele conversion will happen at this marker, once it is included in a gene conversion tract. However, it is difficult to derive a closed form expression for the marginal distribution of *L* when we only allow allele conversions to occur at SNV positions, and with differing rates at each SNV position. Thus, we let allele conversions occur with the same probability *ψ*_*j*_ at all positions within detected gene conversion tract *j*. We use the average heterozygosity rate of positions near detected tract *j* to estimate *ψ*_*j*_.

Letting *a*_*j*_ and *b*_*j*_ (*a*_*j*_ ≤ *b*_*j*_) represent the positions on the chromosome corresponding to the furthest allele converted markers within detected gene conversion tract *j*, we average the heterozygosity rate across the set of positions [*a*_*j*_ − 5000, *b*_*j*_ + 5000] to estimate *ψ*_*j*_:

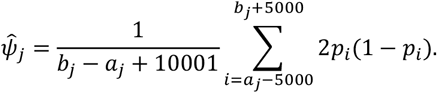

Here, *p*_*i*_ denotes the MAF of position *i* on the chromosome in which the gene conversion event occurred. *p*_*i*_ is calculated using the sample of 125,361 White British individuals from the UK Biobank. Variants with MAF less than 5% were excluded when detecting allele conversions, so we cannot observe allele conversions at these positions (see the section, Detecting gene conversion tracts). Therefore, if the MAF is less than 5% at position *i*, we set *p*_*i*_ = 0. The formula 2*p*(1 − *p*) for heterozygosity at a marker assumes that Hardy-Weinberg equilibrium holds, which is a reasonable approximation for common variants in a relatively homogeneous population.

If either *a*_*j*_ − 5000 or *b*_*j*_ + 5000 exceeds the end of the chromosome, the averaging only takes place within the bounds of the chromosome (e.g. if *a*_*j*_ = 100 and *b*_*j*_ = 200, we only average the heterozygosity rate from positions 1 to 5,200).

### Maximum likelihood estimation of the mean gene conversion tract length

Given observed tract lengths {ℓ_*j*_|*j* = 1, …, *m*}, we propose the following maximum likelihood estimator for *ϕ*, the mean gene conversion tract length, when the gene conversion tract length *N* is drawn from a geometric distribution. Recall that the version of the model in which *N* is geometric was parameterized by λ = 1/*ϕ*, but we can simply maximize with respect to *ϕ*. In other words,

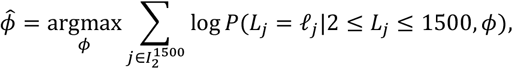

where 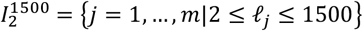. When *N* is a sum of two geometric random variables, we parameterize the distribution of *L* using *γ* = 2/*ϕ* (see Appendix). Unlike the geometric case, our marginal distribution of *L*_*j*_ truncated between 2 and 1,500 still depends on *ψ*_*j*_, so for each *j*, we plug in our estimated 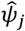 in place of *ψ*_*j*_. Then, we can again maximize with respect to *ϕ*:

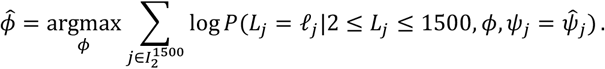

When *N* is a mixture of two geometric components, we have three unknown parameters *ϕ*_1_, *ϕ*_2_, and *w*_1_, which represent the mean of the first component, the mean of the second component, and the mixing proportion of the first component (see Appendix). Again, our marginal distribution of *L*_*j*_ truncated between 2 and 1,500 still depends on *ψ*_*j*_, so for each *j*, we plug in our estimated 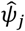 in place of *ψ*_*j*_. Then, we can maximize with respect to *ϕ*_1_, *ϕ*_2_, and *w*_1_:

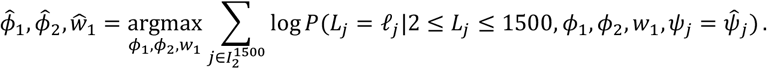

To find the argmax when *N* is geometric or a sum of two geometric random variables, we use the L-BFGS-B algorithm implemented in the scipy.optimize.minimize function from the SciPy Python library.^17^ When *N* is a mixture of two geometric components, we define a grid for *w*_1_ ranging from 0.002 to 0.5, using increments of 0.00025 between 0.002 and 0.01, and increments of 0.05 between 0.05 and 0.5. We chose a finer grid at smaller values of *w*_1_ because preliminary analyses of observed tract lengths from the UK Biobank whole autosome data consistently inferred *w*_1_ to be close to zero. Then, for each *w*_1_ value in the grid, we again ran the L-BFGS-B algorithm from four starting values of (*ϕ*_1_, *ϕ*_2_): (0.0005, 0.0005), (0.0005, 0.1), (0.1, 0.0005), and (0.1, 0.1). Multiple starting values were used because the likelihood of (*ϕ*_1_, *ϕ*_2_) (fixing *w*_1_) appeared to have multiple local maxima. The final maximum likelihood estimates were selected as the set of (*w*_1_, *ϕ*_1_, *ϕ*_2_) values achieving the highest joint likelihood across all grid points of *w*_1_ and starting values of L-BFGS-B.

To choose between the three distributions of *N*, we propose calculating the Akaike Information Criterion (AIC) under each version of the model.^11^ Lower AIC indicates that the distribution of *N* that is used is a better fit to the data.

### Bootstrap confidence intervals

We calculate 95% bootstrap confidence intervals for *ϕ* (*w*_1_, *ϕ*_1_, *ϕ*_2_ in the case where *N* is a mixture of two geometric components). We denote the number of detected gene conversion tracts with observed tract length between 2 and 1,500 bp as 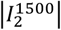. To obtain each bootstrap sample, we sample with replacement 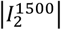 observed tract lengths from the set {ℓ_*j*_|*j* = 1, …, *m*, 2 ≤ ℓ_*j*_ ≤ 1500}. Each bootstrap sample consists of the set of observed tract lengths {ℓ_*j*_} and allele conversion probabilities {*ψ*_*j*_} corresponding to the resampled indices.

We refit our model to 500 bootstrap samples and obtain a new maximum likelihood estimate of *ϕ* (or *w*_1_, *ϕ*_1_, *ϕ*_2_ in the case where *N* is a mixture of two geometric components) for each bootstrap sample. We take the 0.025 and 0.975 quantiles of the resulting bootstrap distributions and use this as the bounds of our 95% bootstrap confidence intervals.

### Simulation study

We use simulated data described in Browning and Browning (2024).^9^ 20 regions of length 10 Mb were generated for 125,000 individuals using the coalescent simulator *msprime* v1.2.^12^ The demographic model for the simulation was an exponentially growing population with an initial size of 10,000 and a growth rate of 3% per generation for the past 200 generations. To simulate recombination and mutation, a crossover rate of 1 cM/Mb and a mutation rate of 1.5 × 10^−8^ per bp per meiosis were used. The mutation rate used is similar to previously inferred mutation rates using IBD segments.^18,19^ Gene conversions were simulated with an initiation rate of 0.02 per Mb and gene conversion lengths were simulated from a geometric distribution with a mean tract length of 300 bp. The processes used to add uncalled deletions and genotype errors are described in Browning and Browning (2024).^9^ Variants with MAF ≤ 0.01 were excluded, the phase information was removed, and Beagle 5.4 was used to statistically phase the genotypes.^14^ The multi-individual IBD analysis detected 284,838 allele conversions belonging to 226,007 detected gene conversion tracts across the 20 regions. We fit our model to the detected gene conversion tracts in each of the 20 regions to estimate the mean gene conversion tract length in each region. For the purposes of this simulation study, we refer to the detected gene conversion tracts in each region as a separate replicate dataset. We refer to fitting our model to the detected gene conversion tracts in each of the 20 regions as a separate replicate of this simulation study.

We fit our model under all three distributions for the true tract length (geometric, sum of two geometric random variables, and mixture of two geometric components). Because the true tract lengths in this simulation study are drawn from a geometric distribution, we are interested in whether the version of the model in which the tract length is geometric will be favored using AIC.

*msprime* only allows gene conversion tract lengths to be drawn from a geometric distribution.^12^ Thus, to test the robustness of our method to different tract length distributions, we run an additional simulation study drawing gene conversion tract lengths from various distributions, including a mixture of two geometric components (see Appendix).

### UK Biobank analysis

We previously described how we obtain the observed tract lengths of all detected gene conversion tracts from the UK Biobank whole autosome data, denoted {ℓ_*j*_|*j* = 1, …, *m*}. We fit our model on this dataset, using all three tract length distributions (geometric, sum of two geometric random variables, and mixture of two geometric components). We further compare model fit under each of these distributions using AIC.

In addition, we run a stratified analysis, stratifying observed tract lengths based on whether they overlapped with a crossover hotspot. To avoid ascertainment bias, where longer tracts are more likely to overlap a crossover hotspot by chance, we defined overlap based on whether the midpoint of the detected gene conversion tract was inside a crossover hotspot. To define crossover hotspots, we use the deCODE genetic map from Halldorsson et al. and follow their definition of crossover hotspots as regions with crossover rates exceeding ten times the genome-wide average.^20^

We calculate local crossover rates between nearby markers on each chromosome by dividing the genetic distance between the two markers by their physical distance. Initially, we calculate the local crossover rate between the first marker in the genetic map, and the marker closest to it that is distant by at least 2 kb. We next calculate the local crossover rate between this newly identified marker and the marker closest to it that is distant by at least 2 kb. We repeat this process until the last marker on this chromosome is included in a local crossover rate calculation, or until we cannot identify further markers that are at least 2 kb away.

If the local crossover rate between two markers is more than ten times the genome-wide average, we classify the region spanning these markers as a crossover hotspot. We stratify the observed tract lengths {ℓ_*j*_|*j* = 1, …, *m*} based on whether the midpoint of the corresponding detected gene conversion tract was inside a crossover hotspot. We then fit our model, separately for each set of tracts. We again use all three tract length distributions to fit the model in this stratified analysis, and compare model fit using AIC.

## Results

### Simulation study

We fit our model to the observed tract lengths from each replicate of the simulation study. The number of observed tract lengths between 2 bp and 1.5 kb across the 20 replicates ranged from 2,005 to 2,314. Recall that a geometric distribution with mean 300 bp was used to simulate gene conversion tract lengths in this simulation study. We estimate the mean tract length under all three tract length distributions (geometric, sum of two geometric random variables, and mixture of two geometric components).

Estimates and confidence intervals using the geometric setting are shown in Figure 1. The average estimate of the mean tract length across the 20 replicates is 289.5 bp under the geometric setting, which is slightly lower than the true mean of 300 bp used to simulate the gene conversion tracts. Under the geometric setting, the true mean of 300 bp is contained in our 95% bootstrap confidence intervals in 14 out of the 20 replicates.

**Figure 1.**
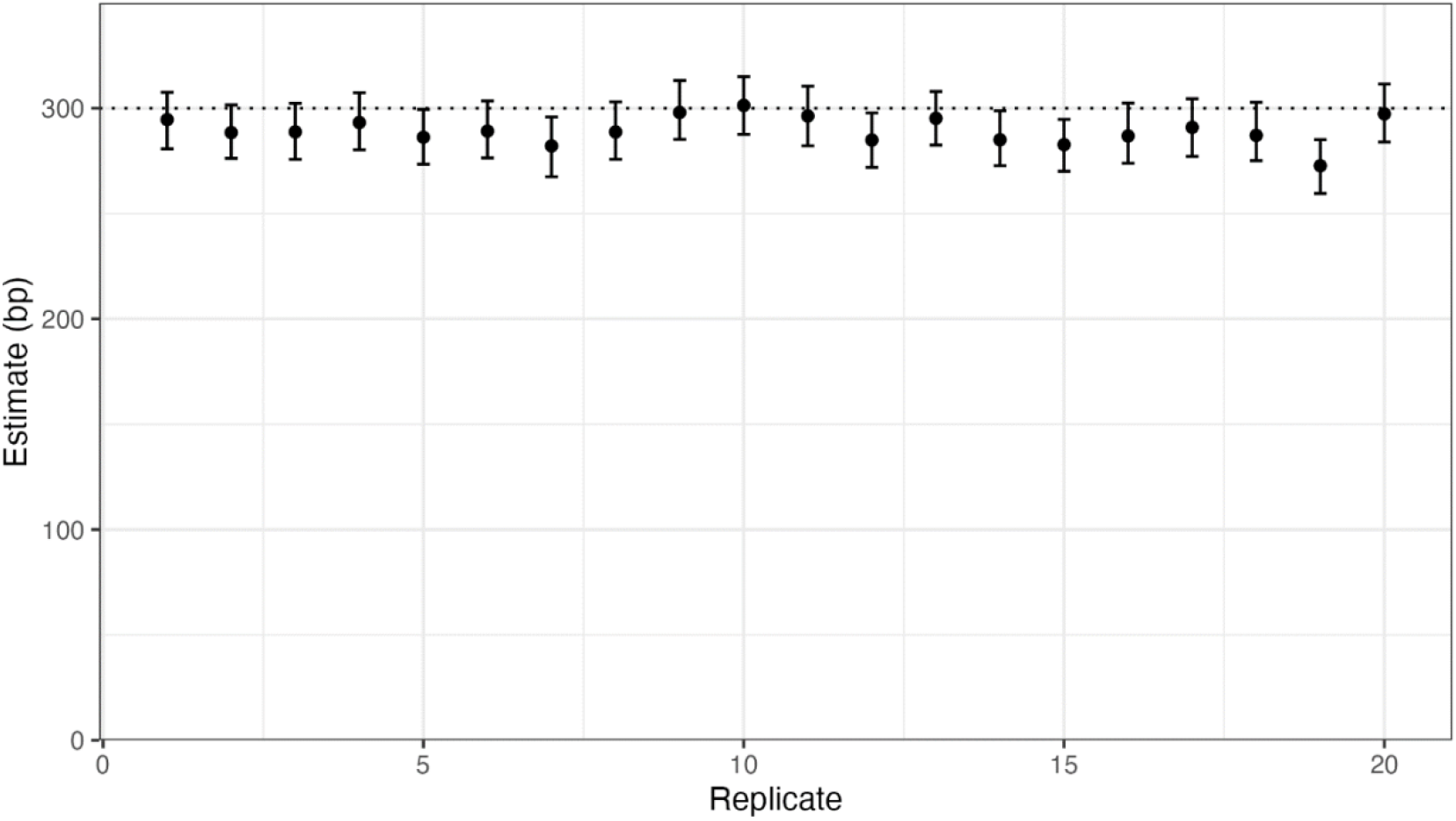
The estimated mean gene conversion tract length under the geometric setting across replicate simulations. The dotted horizontal line represents the true mean gene conversion tract length. Gene conversion tract lengths were simulated using a geometric distribution. We plot our estimate and 95% bootstrap confidence interval under the geometric setting for each replicate simulation.

The geometric setting results in the smallest AIC in 16 out of the 20 replicates. For the remaining four replicates, AIC is lowest when gene conversion tract lengths are assumed to be drawn from a mixture of two geometric components. Estimates for these four replicates using the mixture setting are shown in Figure 2.

**Figure 2.**
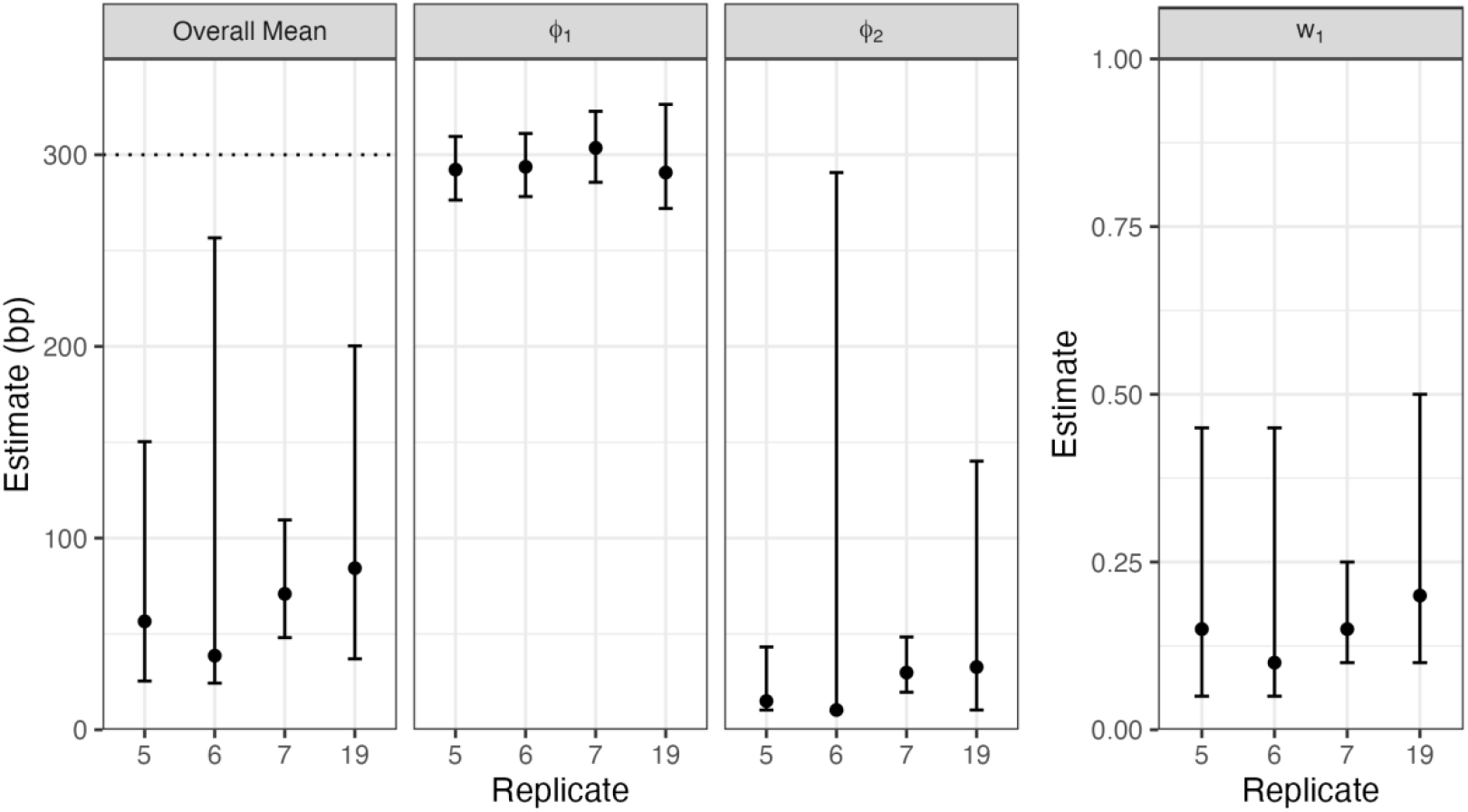
Parameter estimates for four replicates using the mixture distribution. The dotted horizontal line represents the true mean gene conversion tract length. We plot the estimated parameter values with 95% bootstrap confidence intervals for each replicate simulation.

For these four replicates, we see that the mixture setting underestimates the overall mean of 300 bp. Notice that the mean of the first component is estimated to be close to 300 bp for these replicates, but the mean of the second component is estimated to be much lower. The mixing proportion of the first component is estimated to be between 0.1 and 0.2 across the four replicates. 95% confidence intervals for parameters tend to be wide, except for the mean of the first component.

Although the mixture setting results in estimates of the overall mean that are much lower compared to the geometric setting for these four replicates, the difference in AIC between these settings are very small for two of the four replicates (1.6 and 0.8). The difference in AIC for the remaining two replicates are 22.9 and 14.6.

Across all 20 replicates, the difference in AIC between the geometric and mixture settings (positive values preferring the mixture setting) range from 22.9 to –4. An AIC difference of –4 indicates that the log-likelihoods of the two settings were equal, and the difference between the AICs is because of the two additional parameters used under the mixture setting. Because the geometric distribution is nested within the mixture of two geometric components, the log-likelihood under the geometric setting cannot exceed that of the mixture setting.

### UK Biobank analysis

We applied our estimation method to the observed tract lengths detected from the UK Biobank whole autosome data. The AIC is lowest (indicating best fit) under the setting where the true tract length distribution is assumed to be a mixture of two geometric components (11,860,323). The AIC for the geometric and sum of two geometric settings were 12,201,916 and 12,268,153 respectively. The difference in AIC between the mixture setting and the geometric setting, which had the next lowest AIC, was 341,593, providing strong evidence in favor of the mixture setting.

When assuming that gene conversion tract lengths are a mixture of two geometric components, we estimate the mixing proportion for the first component to be 0.00525 (95% CI: [0.005, 0.00525]). We estimate the mean of the first and second components to be 724.7 bp (95% CI: [720.1, 728.7]) and 16.9 bp (95% CI: [16.4, 17.0]) respectively. We estimate the overall mean to be 20.6 bp (95% CI: [19.9, 20.7]).

For the stratified analysis, we calculated the genome-wide average crossover rate to be 1.23 cM/Mb. We classify any regions exceeding ten times this rate as a crossover hotspot. Of the 876,584 tracts detected from the UK Biobank sequence data, the midpoints of 290,766 (33.2%) were contained within a crossover hotspot. For both tract sets, the set of tracts with midpoint in a crossover hotspot and the remaining tracts, the lowest AIC was obtained under the mixture setting, so we report our results from assuming that gene conversion tract lengths are drawn from the mixture distribution.

For detected tracts with midpoint located within a crossover hotspot, we estimate the mean of the first and second components to be 579.8 bp (95% CI: [574.8, 585.5]) and 20.3 bp (95% CI: [19.7, 21.1]) respectively. We further estimate the mixing proportion for the first component to be 0.0095 (95% CI: [0.00925, 0.01]). We estimate the overall mean to be 25.6 bp (95% CI: [24.9, 26.7]).

For detected tracts with midpoint not located within a crossover hotspot, we estimate the mean of the first and second components to be 813.9 bp (95% CI: [807.7, 819.3]) and 15.5 bp (95% CI: [14.9, 15.6]) respectively. We further estimate the mixing proportion for the first component to be 0.004 (95% CI: [0.00375, 0.004]). We estimate the overall mean to be 18.7 bp (95% CI: [17.9, 18.8]).

### Discussion

Previous studies have tried to measure gene conversion tract lengths in humans by detecting allele conversions from pedigree and sperm-typing data.^1,3–5^ However, in these studies, it is only possible to detect gene conversion events occurring in a relatively small number of meioses. Efforts to detect gene conversions from pedigree data have been limited by the number of multi-generational pedigrees that have been genotyped. Sperm-typing studies have also been limited by the availability of appropriate data. In sperm-typing studies, distinguishing genotype errors from allele conversions is also difficult.

By applying the multi-individual IBD method to the UK Biobank whole autosome data, we were able to detect gene conversion events across multiple meioses in the ancestral history of this population.^9^ Using this method, 5,961,128 gene conversion tracts were detected, which is several orders of magnitude larger than what had been detected in humans in the past. In the largest pedigree study conducted to detect gene conversions, less than 30,000 gene conversion events were detected from 5,420 trios.^8^

We proposed a likelihood-based estimation method to infer the mean gene conversion tract length. Our method is inspired by a previous approach developed by Betran et al., which was applied to gene conversion tracts detected in 34 *Drosophila subobscura* sequences.^16^ However, we made several key improvements. First, we define a separate allele conversion probability for each gene conversion tract, based on the density and heterozygosity rate of markers near each tract. Second, we allow gene conversion tract lengths to follow multiple distributions, including a mixture of two geometric components, which has previously been found to appropriately model gene conversion tract lengths in other mammals.^6^ Third, we derive the closed-form expression for the distribution of observed tract lengths for each true tract length distribution, which allows for fast and exact calculation of the joint likelihood during maximum likelihood estimation. Finally, we allow for the selection of the best fitting tract length distribution using AIC.

We ran a coalescent simulation incorporating gene conversion events to validate our estimation method. Since we used *msprime* for the simulation, gene conversion tract lengths were necessarily drawn from a geometric distribution. Nonetheless, this simulation allowed us to accurately capture potential biases arising from evolutionary and technical factors such as mutations and genotype errors, as well as potential artifacts introduced by the multi-individual IBD detection method used to identify gene conversion tracts.^9^ We found that our model accurately estimated the mean gene conversion tract length when the length distribution of gene conversion tracts was correctly specified to be geometric. Our model resulted in biased estimates of the mean gene conversion tract length when the length distribution was incorrectly specified. In most replicates of this simulation study (16 out of 20 replicates), AIC correctly determined the best fitting distribution to be geometric.

To further assess the robustness of our model to the misspecification of the tract length distribution, we conducted a separate simulation study where gene conversion tract lengths were drawn from multiple distributions (see Appendix). In this study, we found that the model selected by AIC consistently produced relatively unbiased estimates across a range of tract length distributions. Furthermore, when the true tract length distribution was one of the three distributions that we allow for in our model, we found that AIC selects the true distribution in most cases.

Applying our method to observed tract lengths detected from the UK Biobank whole autosome data, we found that the mixture setting, which had the lowest AIC by a large margin, estimated most tracts have a small mean of 16.9 (95% CI: [16.4, 17.0]), and only a small proportion of tracts have a much larger mean of 724.7 (95% CI: [720.1, 728.7]). The mixing proportion for the geometric distribution with the smaller mean was estimated to be 0.00525 (95% CI: [0.005, 0.00525]). We estimate the overall mean to be 20.6 (95% CI: [19.9, 20.7]).

Our estimate of the mean gene conversion tract length is very sensitive to the assumed tract length distribution. When assuming that gene conversion tract lengths are geometric, our model estimates the mean gene conversion tract length to be 459.0 bp (95% CI: [457.3, 460.5]), which is much higher than our estimate under the mixture setting. However, given the large AIC difference between these two models (341,593), we are confident that the mixture distribution is a much better fit to the data. This result aligns with previous findings in humans. Palsson et al. found that among tracts shorter than 1 kb, the majority had a smaller mean length compared to the longer tracts.^8^ The higher estimate we obtained under the geometric distribution is also consistent with our simulation results. In the simulation assessing the robustness of our method, where we draw gene conversion tract lengths from various distributions (see Appendix), we found that assuming a geometric distribution when the true distribution is a mixture of two geometric components can lead to an inflated estimate of the mean tract length, particularly when one component has a substantially larger mean but contributes relatively few tracts (see Table 1).

**Table 1.** Results from simulation study to assess robustness. We assess the performance of our method under each distribution that we use to simulate the true tract lengths (first column) and the chosen setting of the tract length distribution (second column). We report the empirical bias (third column) and standard deviation (fourth column) of our estimate of the mean, as well as the empirical coverage of our 95% confidence interval (fifth column) across 100 replicates of the simulation study. Under the AIC-selected setting, we use the estimate and confidence interval from the distributional setting with the smallest Akaike Information Criterion (AIC) value in each of the 100 replicates.

| Distribution | Chosen Setting | Bias | SD | Coverage |
| --- | --- | --- | --- | --- |
| Geom | AIC-selected | 13.9 | 29.2 | 0.54 |
|  | Mixture | -21.1 | 26.8 | 0.79 |
|  | Geom | -2.5 | 7.9 | 0.85 |
|  | Geom2 | 45.9 | 11.8 | 0.01 |
| Geom2 | AIC-selected | -5.0 | 13.9 | 0.79 |
|  | Mixture | -34.8 | 7.7 | 0.00 |
|  | Geom | -33.3 | 4.5 | 0.00 |
|  | Geom2 | -0.5 | 6.6 | 0.93 |
| Geom3 | AIC-selected | -18.4 | 8.3 | 0.12 |
|  | Mixture | -45.1 | 5.8 | 0.00 |
|  | Geom | -43.9 | 3.3 | 0.00 |
|  | Geom2 | -16.4 | 4.9 | 0.15 |
| Mixture | AIC-selected | 7.3 | 27.0 | 0.98 |
|  | Mixture | 7.3 | 27.0 | 0.98 |
|  | Geom | 265.6 | 34.6 | 0.00 |
|  | Geom2 | 434.8 | 45.6 | 0.00 |
| Uniform | AIC-selected | -23.6 | 4.4 | 0.00 |
|  | Mixture | -48.3 | 3.1 | 0.00 |
|  | Geom | -48.3 | 3.1 | 0.00 |
|  | Geom2 | -23.6 | 4.4 | 0.00 |

We estimated the overall mean gene conversion tract length to be 20.6 bp (95% CI: [19.9, 20.7]), which is shorter than previous estimates. For instance, Palsson et al. reported mean tract lengths of 123 bp (95% CI: [94, 135]) for paternal and 102 bp (95% CI: [71, 125]) for maternal transmissions.^8^ Methodological differences between our approach and the NCOurd model used by Palsson et al. may account for this discrepancy.^7^ NCOurd requires specifying a penetrance parameter, defined as the probability that a heterozygous marker within a gene conversion tract is allele converted. In our framework, we set the allele conversion probability within each tract equal to the local mean heterozygosity rate. This effectively assumes that, for shorter gene conversion tracts (<1.5 kb), all heterozygous markers are allele converted. This would correspond to using a penetrance of one in NCOurd. In contrast, Palsson et al. estimate a fixed penetrance of 0.66 for all detected tracts by using a grid of penetrance values and selecting the one that maximizes the model likelihood. This implies that roughly a third of heterozygous sites within a gene conversion tract do not undergo allele conversion, leading to longer estimated tract lengths. Importantly, penetrance may vary with tract length, making the use of a single penetrance value potentially inappropriate. However, estimating penetrance as a function of the tract length is challenging, especially for short tracts, which often do not overlap with many markers. This limitation has been noted in the original NCOurd publication.^7^

There are a few other findings on the length distribution of gene conversion tracts in humans, most notably, in the sperm-typing study by Jeffreys and May, which concluded that the mean length is in the range of 55-290 bp.^3^ Jeffreys and May inferred the range of mean gene conversion tract lengths (55-290 bp) by comparing observed gene conversion lengths to simulated tracts under geometrically and normally distributed gene conversion tract lengths. However, our simulation where tract lengths are drawn from a mixture distribution suggests that modeling all tracts using a single distribution, without explicitly accounting for outliers, can lead to an inflated estimate of the mean when a small proportion of tracts are much longer than the rest (see Appendix).

Wall et al. analyzed gene conversion tracts shorter than 10 kb in a captive baboon colony using a mixture of two geometric distributions. They estimated that 99.8% of tracts had a mean length of 24 bp (95% CI: [18, 31]), while the remaining tracts had a mean of 4.3 kb (95% CI: [2.6, 4.9]). Both the mixing proportion and the mean of the shorter component are similar to our estimates.

We ran an additional analysis in which we stratified detected gene conversion tracts from the UK Biobank whole autosome data by whether their midpoints were located within a crossover hotspot. In both sets of tracts, the set of tracts with midpoints located within a crossover hotspot and the remaining tracts, AIC was smallest when assuming a mixture distribution for the true tract length distribution. Comparing the estimated parameters for the mixture distribution in each set, detected tracts with midpoints located within a hotspot were estimated to have a larger proportion of longer tracts (0.0095; 95% CI: [0.00925, 0.01]) compared to the remaining detected tracts (0.004; 95% CI: [0.00375, 0.004]). The mean of the longer component of the mixture distribution was estimated to be smaller for hotspot tracts (579.8 bp; 95% CI: [574.8, 585.5]) compared to the remaining tracts (813.9 bp; 95% CI: [807.7, 819.3]). The mean of the shorter component of the mixture distribution was estimated to be larger for hotspot tracts (20.3 bp; 95% CI: [19.7, 21.1]) compared to the remaining tracts (15.5 bp; 95% CI: [14.9, 15.6]). The overall mean was larger for hotspot tracts (25.6 bp; 95% CI: [24.9, 26.7]) compared to the remaining tracts (18.7 bp; 95% CI: [17.9, 18.8]). These differences in the proportion of longer tracts, and in the mean lengths of the shorter and longer components were significant. This is a preliminary finding and we recommend further analysis to confirm this result. Recombination hotspots correlate with other genomic features such as GC rate,^21^ so the difference may be caused by factors other than the recombination rate itself.

It is important to acknowledge that our method omits observed tract lengths exceeding 1.5 kb, because we cannot accurately detect observed tract lengths corresponding to longer gene conversion tracts. Complex gene conversion events, which result in both allele converted and non-allele converted markers, often span more than 1.5 kb.^5^ To appropriately model the lengths of these longer tracts, we would need to apply a detection method that can reliably detect these tracts.

In this study, we did not extend the mixture distribution, which was strongly favored by AIC, to have more than two components. While a mixture model with additional components may better capture the true distribution of gene conversion tract lengths, exploring such models proved computationally challenging due to the complexity of the optimization procedure and the large number of detected gene conversion tracts. Future work may consider more flexible models, such as three-component mixtures, particularly as methods for detecting longer or complex gene conversion events from population-level sequence data become available.

## Appendix

### Deriving the marginal distribution of the observed tract length under two alternative settings

We first consider the case in which *N* is distributed as a sum of two independent and identically distributed geometric random variables each with mean *ϕ*/2. We have,

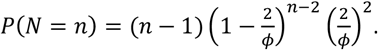

Letting 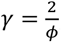,

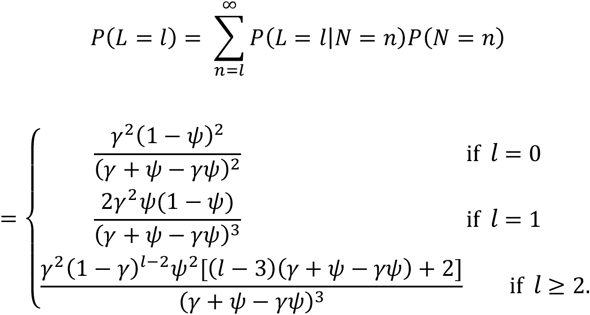

Then,

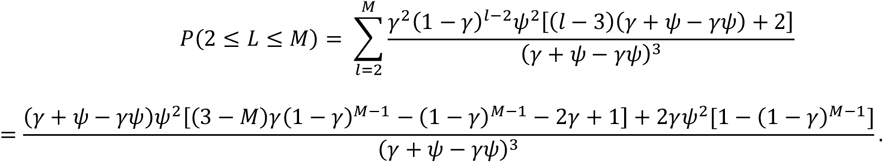

Then,

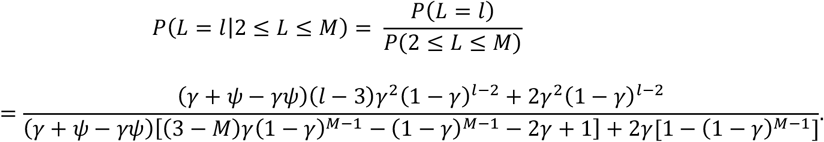

Similarly to the case where *N* is geometric, we index our random variable *L* using *j* so that *L*_*j*_ represents the random variable corresponding to the observed tract length for detected tract *j* in our dataset. This time, we also index *ψ* using *j* so that an allele conversion happens with probability *ψ*_*j*_ at every position within the *j* th detected tract (the estimation of *ψ*_*j*_ is described in the section, Estimating the allele conversion probability for each detected tract). We have,

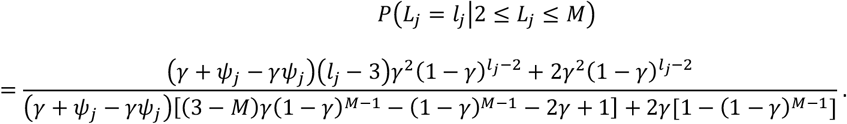

We next consider the case where *N* is distributed as a mixture of two geometric components. We let the two geometric means be *ϕ*_1_ and *ϕ*_2_, and let *w*_1_ represent the mixing proportion of the first component. We have,

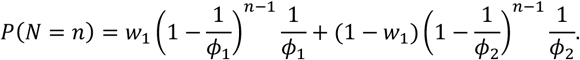

Letting λ_1_ = 1/*ϕ*_1_ and λ_2_ = 1/*ϕ*_2_,

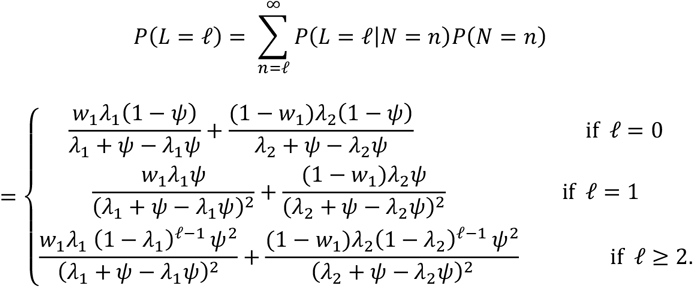

Then,

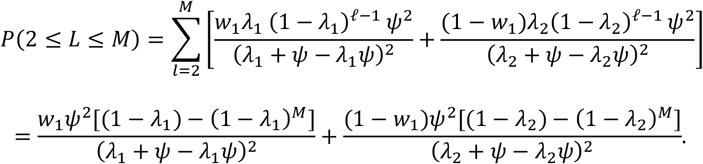

Then,

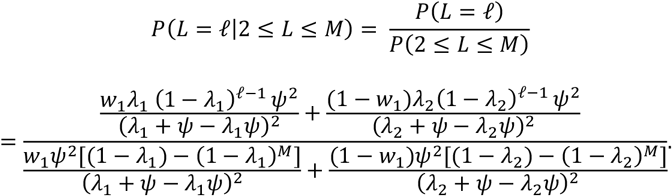

Again using *j* to index detected tracts,

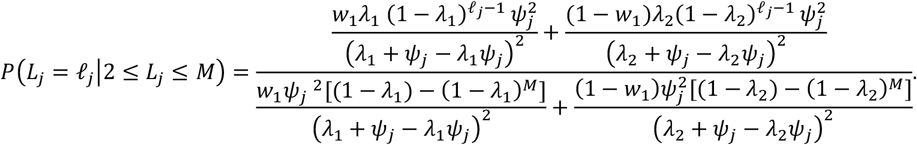

In practice, we plug in *M* = 1500 because we exclude all observed tract lengths longer than 1500 bp detected from the UK Biobank whole autosome data.

### Simulation study to assess the robustness of the model

We run a simulation study to assess how well our model can estimate the mean tract length *ϕ* when gene conversion tract lengths are from various distributions. We simulate observed tract lengths {ℓ_*j*_|*j* = 1, …, *m*} using five distributions for the length distribution of gene conversion tracts (Figure S1):

1. Geometric distribution with mean 100 bp
2. Sum of two geometric random variables, each with mean 50 bp
3. Sum of three geometric random variables, each with mean 33.3 bp
4. Discrete uniform distribution with support from 1 to 199 bp
5. Mixture of two geometric components with means 700 bp and 68.4 bp, with 5% of tracts being drawn from the first component

All five distributions have an overall mean of 100 bp. Recall that in the previous coalescent simulation, we generated 20 regions of length 10 Mb for 125,000 individuals using the coalescent simulator *msprime* v1.2.^12^ In this simulation study, we generate observed tract lengths by simulating gene conversion tracts on the first region (out of the 20 regions) from the previous coalescent simulation. To simulate one set of observed tract lengths, we first sample 100,000 individuals with replacement from the 125,000 individuals. For each resampled individual, we follow these steps:

1. We randomly select a starting position for the gene conversion tract, chosen uniformly across the 10 Mb region.
2. We draw the length of the gene conversion tract from one of the five specified distributions.
3. We determine the observed tract length as the length spanning the furthest heterozygous markers within the simulated gene conversion tract.

This procedure results in 100,000 observed tract lengths, some of which may be zero bp due to the absence of heterozygous markers within the corresponding gene conversion tracts. For each of the five distributions listed earlier, we repeat this procedure 100 times to obtain 100 sets of observed tract lengths. Then, we fit our model under all distributions of the true tract length (geometric, sum of two geometric random variables, and a mixture of two geometric components) to each set of observed tract lengths. Because the number of observed tract lengths differ for each set, we sample 200 observed tract lengths between 2 and 1,500 bp in each set to make sure that we use the same number of observed tract lengths for estimation.

For each set of observed tract lengths, and for each assumed distribution for the true tract length, we obtain both a point estimate and a 95% bootstrap confidence interval for the mean tract length. Table 1 reports the empirical bias and empirical standard deviation of our estimate of the mean, as well as the empirical coverage probability of our 95% confidence interval under all model settings across 100 sets of observed tract lengths generated using each of the five distributions. Under the AIC-selected setting, we use the estimate and confidence interval from the assumed tract length distribution with the smallest AIC value in each set of observed tract lengths. Table 2 reports the number of times each assumed tract length distribution was preferred by AIC, across the 100 sets of observed tract lengths generated using each of the five distributions.

**Table 2.** Number of replicates each distributional setting was selected by the Akaike Information Criterion (AIC). For each of the five data-generating distributions, we simulated 100 sets of observed tract lengths. We then counted how many times each distribution of *N* was selected as the best fitting distribution based on AIC.

| Distribution | Chosen Setting | Times Selected by AIC |
| --- | --- | --- |
| Geom | Mixture | 3 |
|  | Geom | 59 |
|  | Geom2 | 38 |
| Geom2 | Mixture | 0 |
|  | Geom | 14 |
|  | Geom2 | 86 |
| Geom3 | Mixture | 0 |
|  | Geom | 7 |
|  | Geom2 | 93 |
| Mixture | Mixture | 100 |
|  | Geom | 0 |
|  | Geom2 | 0 |
| Uniform | Mixture | 0 |
|  | Geom | 0 |
|  | Geom2 | 100 |

## Acknowledgements

This research has been conducted using the UK Biobank Resource under application number 19934. The methodological and analytical work performed in this study was supported by the National Human Genome Research Institute (NHGRI) under award number R01 HG005701. The content is solely the responsibility of the authors and does not necessarily represent the official views of the National Institutes of Health or the UK Biobank.

## Data and code availability

The data and the code generated during this study are available at https://github.com/nobuakimasaki/gene-conversion-lengths. This includes the full list of positions for the 5,961,128 gene conversion tracts detected from the UK Biobank whole autosome sequence data.

