## Supplementary Materials for "Modeling the length distribution of gene conversion tracts in humans from the UK Biobank sequence data"

### Supplemental figures

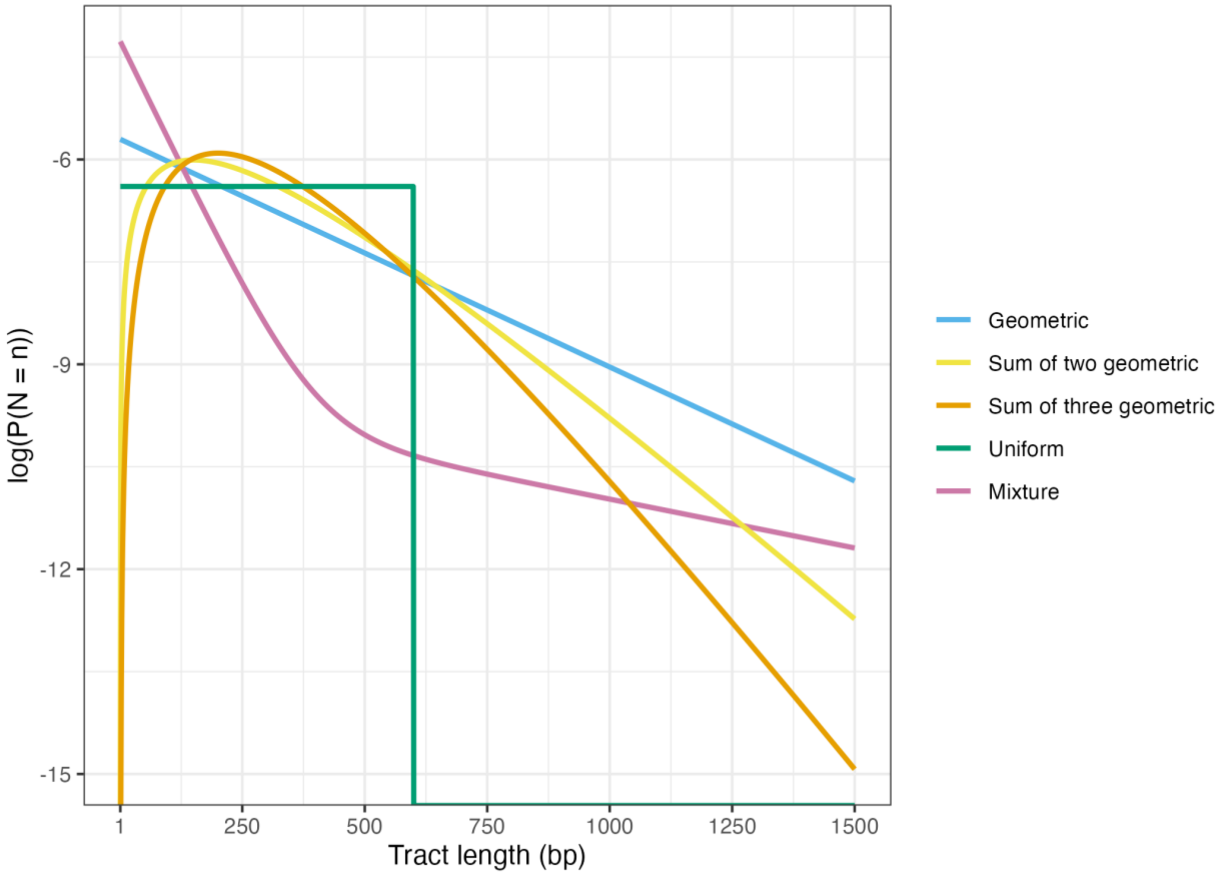


**Figure S1. Probability distribution functions (log scale) of the five distributions used to simulate gene conversion tract lengths.** We plot the distribution functions of the geometric distribution, the sum of two geometric random variables, the sum of three geometric random variables, the discrete uniform distribution, and the mixture of two geometric components that we draw the gene conversion tract lengths from the simulation study used to assess the robustness of the model.

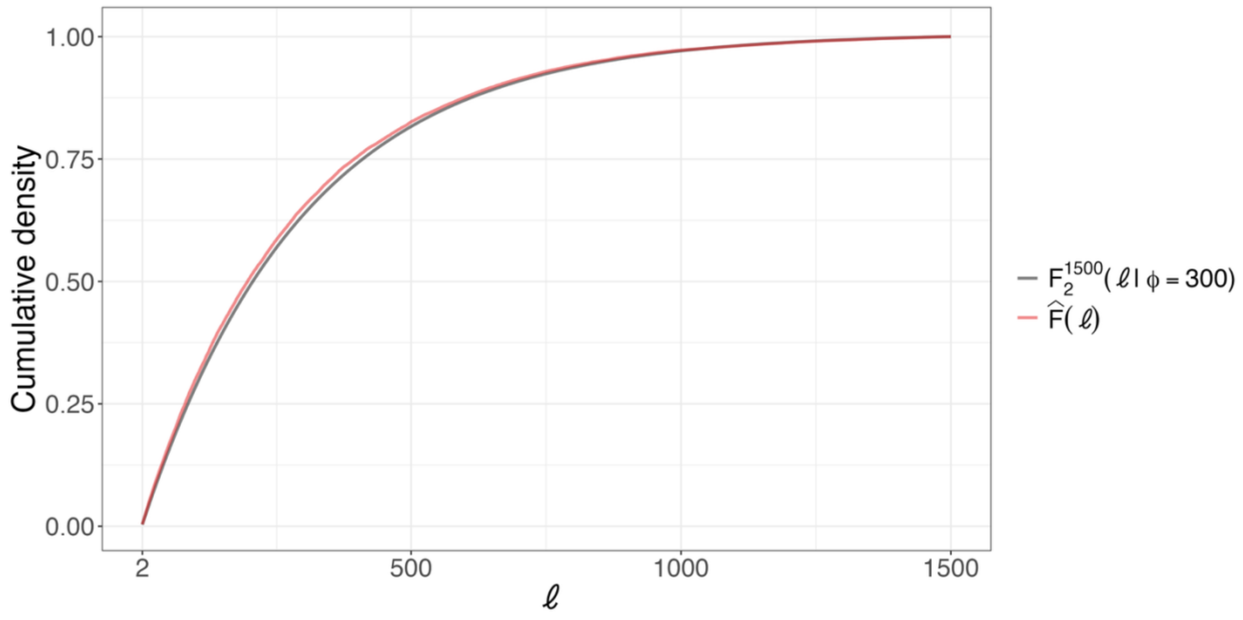


**Figure S2. Comparing the CDF of $L$ and the empirical CDF of observed tract lengths detected in the coalescent simulation.** We plot the CDF of $L$ truncated between 2 and 1,500 bp (in grey) and the empirical CDF of observed tract lengths between 2 and 1,500 bp detected in the coalescent simulation (in red).

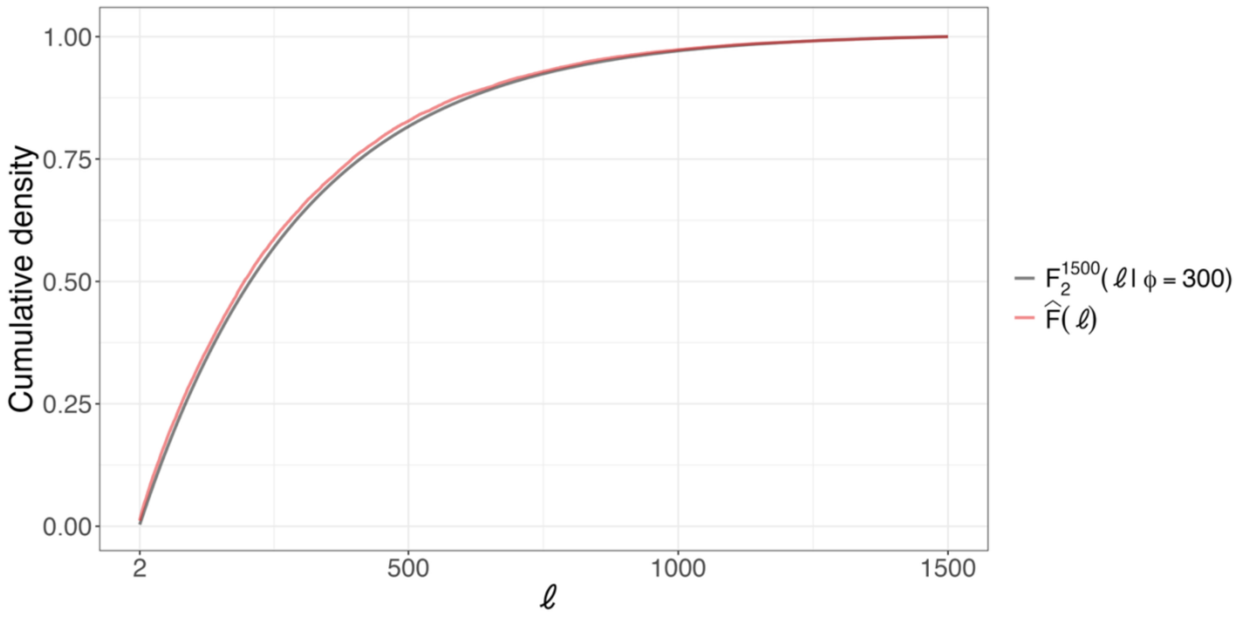


**Figure S3. Comparing the CDF of $L$ and the empirical CDF of observed tract lengths generated in the simulation without linkage disequilibrium.** We plot the CDF of $L$ truncated between 2 and 1,500 bp (in grey) and the empirical CDF of observed tract lengths between 2 and 1,500 bp generated in the simulation without linkage disequilibrium (in red).

### Supplemental text

#### Text S1

In this section, we specify gene conversion tract lengths to be geometric. Recall that $\phi$ is the mean gene conversion tract length. Then, the observed tract length distribution for detected gene conversion tract $j$, truncated between 1 and 1,500 bp, is,

$$P(L_j = \ell_j | 1 \leq L_j \leq 1500, \lambda, \psi_j) = \begin{cases} \dfrac{\lambda\psi_j}{\lambda\psi_j + \psi_j^2[1-\lambda-(1-\lambda)^{1500}]} & \text{if } \ell_j = 1 \\ \dfrac{\lambda(1-\lambda)^{\ell_j - 1}\psi_j^2}{\lambda\psi_j + \psi_j^2[1-\lambda-(1-\lambda)^{1500}]} & \text{if } \ell_j \geq 2, \end{cases}$$

where $\lambda = 1/\phi$ and $\psi_j$ is the allele conversion probability for detected tract $j$.

In the main text, we described a method for obtaining $\hat{\psi}_j$, our estimate of $\psi_j$, for all detected tracts $j$. Plugging in $\psi_j = \hat{\psi}_j$, we can obtain the probability mass at 1 bp, conditioned on $1 \leq L_j \leq 1500$ and $\lambda$:

$$P(L_j = 1 | 1 \leq L_j \leq 1500, \lambda, \psi_j = \hat{\psi}_j) = \frac{\lambda\hat{\psi}_j}{\lambda\hat{\psi}_j + \hat{\psi}_j^2[1-\lambda-(1-\lambda)^{1500}]}.$$

We can estimate the proportion of detected tracts with an observed tract length of 1 bp (among detected tracts with an observed tract length less than or equal to 1,500 bp) by taking the mean of $P(L_j = 1 | 1 \leq L_j \leq 1500, \lambda, \psi_j = \hat{\psi}_j)$ across all detected tracts $j$ with an observed tract length that is less than or equal to 1,500 bp. Let $\hat{\pi}(L = 1 | 1 \leq L \leq 1500, \lambda)$ denote this estimated proportion, computed as:

$$\hat{\pi}(L = 1 | 1 \leq L \leq 1500, \lambda) = \frac{1}{|I_1^{1500}|} \sum_{j \in I_1^{1500}} P(L_j = 1 | 1 \leq L_j \leq 1500, \lambda, \psi_j = \hat{\psi}_j),$$

where $I_1^{1500} = \{j = 1, \ldots, m | 1 \leq \ell_j \leq 1500\}$ and $|I_1^{1500}|$ represents the number of detected tracts with an observed tract length that is less than or equal to 1,500 bp. In this summation, the only quantity that varies across tracts is $\hat{\psi}_j$, the tract-specific estimate of the allele conversion probability. Notice how $\hat{\pi}(L = 1|1 \leq L \leq 1500, \lambda)$ depends on $\lambda = 1/\phi$, for which we can plug in an appropriate value (an estimate or the true value if it is known).

Once we obtain the observed tract lengths of detected gene conversion tracts, denoted $\{\ell_j | j = 1, \ldots, m\}$, using the multi-individual IBD method,[1] we know the proportion of detected tracts with an observed tract length of 1 bp (among detected tracts with an observed tract length less than or equal to 1,500 bp) in our dataset. If our estimate $\hat{\pi}(L = 1|1 \leq L \leq 1500, \lambda)$ differs from this proportion, our model may not be fitting well to the data.

Browning and Browning ran a coalescent simulation incorporating gene conversions, where they fixed the mean gene conversion tract length to be 300 bp.[1] 20 regions of length 10 Mb were generated for 125,000 individuals, and the multi-individual IBD analysis detected 284,838 allele conversions belonging to 226,007 detected gene conversion tracts across the 20 regions. This simulation is described in more detail in the main text and in Browning and Browning (2024).[1]

From this simulation study, the actual proportion of detected tracts with an observed tract length of 1 bp (among detected tracts with an observed tract length less than or equal to 1,500 bp) was 0.807. However, $\hat{\pi}(L = 1|1 \leq L \leq 1500, \phi = 300) = 0.860$ . This indicates that our model is overestimating the proportion of detected tracts with an observed tract length of 1 bp in the coalescent simulation.

We can similarly compare the actual proportion of detected tracts with an observed tract length of 2 bp or longer to the distribution $P(L = \ell|2 \leq L \leq 1500, \phi)$ derived in the main text. This time, we do not have to worry about varying $\hat{\psi}_j$ for each tract $j$, because our distribution conditional on $2 \leq L \leq 1500$ no longer depends on the allele conversion probability. Thus, we do not have to average across detected tracts like we did earlier. For example, we can compare the actual proportion of detected tracts with an observed tract length of 3 bp (among detected tracts with an observed tract length between 2 and 1,500 bp) to $P(L = 3|2 \leq L \leq 1500, \phi = 300)$. To facilitate this comparison, we define the CDF of $L$ truncated between 2 and 1,500 bp as $F_2^{1500}(\ell|\phi) = \sum_{k=2}^{\ell} P(L = k|2 \leq L \leq 1500, \phi)$. In Figure S2, we plot this and the empirical CDF of observed tract lengths between 2 and 1,500 bp detected in the coalescent simulation. We see from Figure S2 that our truncated distribution of $L$ fits well to the actual proportion of observed tract lengths between 2 and 1,500 bp.

We want to figure out why our model is not fitting well to the actual proportion of detected tracts with an observed tract length of 1 bp in the coalescent simulation. We think this is likely because our model does not account for linkage disequilibrium, even though linkage disequilibrium is present in the simulated regions.

Our model assumes that all positions within a gene conversion tract have the same probability of allele conversion. This means that an allele conversion occurring at one position does not make it more or less likely that an allele conversion will occur at another nearby position within the same gene conversion tract. This assumption is used to derive the marginal distribution of $L$ in the main text. However, in this coalescent simulation and in real populations, linkage disequilibrium can cause heterozygosity to be correlated between nearby positions, leading to allele conversions occurring together at nearby positions more frequently than if these positions were independent from one another. This may explain why the actual proportion of detected tracts with an observed tract length of 1 bp in the coalescent simulation is smaller than what the model predicts.

To test whether linkage disequilibrium is causing a smaller proportion of detected tracts to have an observed tract length of 1 bp compared to what the model predicts, we simulate observed tract lengths under a setting where markers are independent. For this simulation, we use the population heterozygosity rates of markers on chromosome 1 from the UK Biobank whole autosome data. We use the following steps to simulate observed tract lengths:

1. We generate $10^6$ gene conversion tracts by uniformly sampling the starting position on chromosome 1 and drawing the length of the gene conversion tract from a geometric distribution with mean 300. The start and end positions of each tract are saved.
2. We let an allele conversion occur at each position $i$ within each gene conversion tract with probability $2p_i(1-p_i)$, where $p_i$ is the minor allele frequency at position $i$.
3. For each gene conversion tract, we obtain the observed tract length of the gene conversion tract by taking the length spanning the furthest allele converted positions.

In step 2, we set $p_i = 0$ if the minor allele frequency is less than 5% at position $i$ to prevent detecting allele conversions at these markers, like in the multi-individual IBD method.[1]

From this simulation, the actual proportion of observed tract lengths that were 1 bp was 0.813 (among detected tracts with an observed tract length between 1 and 1,500 bp) whereas $\hat{\pi}(L = 1|1 \leq L \leq 1500, \phi = 300) = 0.818$. This time, our model only slightly overestimates this proportion. From Figure S3, we also see that our model closely fits the empirical distribution of observed tract lengths between 2 and 1,500 bp generated from this simulation.

Compared to the coalescent simulation, our model better predicts the proportion of observed tract lengths that are 1 bp from this simulation, in which observed tract lengths are generated under a setting where markers are independent. Recall that in the coalescent simulation, the model overestimates the proportion of observed tract lengths that are 1 bp. This indicates that linkage disequilibrium may cause the proportion of observed tract lengths that are 1 bp to be lower than what the model predicts. When estimating the mean length of gene conversion tracts, we can avoid this issue by only considering observed tract lengths between 2 and 1,500 bp and by truncating the marginal distribution of $L$ between 2 and 1,500 bp before model fitting, as we have done in the main paper.
